# The nature and origin of synaptic inputs to vestibulospinal neurons in the larval zebrafish

**DOI:** 10.1101/2023.03.15.532859

**Authors:** Kyla R. Hamling, Katherine Harmon, David Schoppik

## Abstract

Vestibulospinal neurons integrate sensed imbalance to regulate postural reflexes. As an evolutionarily-conserved neural population, understanding their synaptic and circuit-level properties can offer insight into vertebrate antigravity reflexes. Motivated by recent work, we set out to verify and extend the characterization of vestibulospinal neurons in the larval zebrafish. Using current clamp recordings together with stimulation, we observed that larval zebrafish vestibulospinal neurons are silent at rest, yet capable of sustained spiking following depolarization. Neurons responded systematically to a vestibular stimulus (translation in the dark); responses were abolished after chronic or acute loss of the utricular otolith. Voltage clamp recordings at rest revealed strong excitatory inputs with a characteristic multimodal distribution of amplitudes, as well as strong inhibitory inputs. Excitatory inputs within a particular mode (amplitude range) routinely violated refractory period criteria and exhibited complex sensory tuning, suggesting a non-unitary origin. Next, using a unilateral loss-of-function approach, we characterized the source of vestibular inputs to vestibulospinal neurons from each ear. We observed systematic loss of high-amplitude excitatory inputs after utricular lesions ipsilateral, but not contralateral to the recorded vestibulospinal neuron. In contrast, while some neurons had decreased inhibitory inputs after either ipsilateral or contralateral lesions, there were no systematic changes across the population of recorded neurons. We conclude that imbalance sensed by the utricular otolith shapes the responses of larval zebrafish vestibulospinal neurons through both excitatory and inhibitory inputs. Our findings expand our understanding of how a vertebrate model, the larval zebrafish, might use vestibulospinal input to stabilize posture. More broadly, when compared to recordings in other vertebrates, our data speak to conserved origins of vestibulospinal synaptic input.

## INTRODUCTION

Vestibular reflexes maintain posture in the face of gravity^1^. These reflexes originate with evolutionarily ancient “vestibulospinal” circuits that link the vestibular sensory periphery and the spinal cord^2,3^. Vestibulospinal neurons, first identified by Deiters^4^, are descending projection neurons found in the lateral vestibular nucleus of the hindbrain. Defining the properties of synaptic inputs to vestibulospinal neurons is critical to understand how sensed imbalance is transformed into corrective behaviors.

The larval zebrafish has emerged as a useful model for studying balance behaviors^5^, particularly those mediated by vestibular circuits^6–13^. At 4 days post-fertilization (dpf), larval zebrafish maintain a dorsal-up stable roll posture to navigate in the water column, find food, and avoid predators. To do so, loss-of-function experiments^14–16^ suggest they rely on a sense of gravity mediated by an otolithic (utricular) organ; while present and capable of transduction^17^, their semicircular canals are too small to function under normal conditions^18^. Larval zebrafish are genetically-tractable and largely transparent, allowing for rapid and reliable identification of vestibulospinal neurons^19,20^. Unlike most other preparations, larval zebrafish vestibulospinal neurons are accessible for *in vivo* patchclamp recording allowing characterization of their synaptic inputs.

Recent work^20–22^ suggests a model for how synaptic inputs onto vestibulospinal neurons might shape their response. Vestibular neuron responses are largely linear – a feature thought to facilitate proportional and continuous reflexive responses to destabilization (though see^23^). Unlike most sensory synapses that either adapt or facilitate, the synapse between peripheral afferents and brainstem vestibular neurons has a number of specializations, identified with slice electrophysiology and electron micrography, to allow linear transmission^24,25^. These specializations predict that *in vivo* release from individual vestibular afferents might produce depolarization with a characteristic amplitude in a target vestibulospinal neuron. Recent findings by Liu et. al support this model: excitatory synaptic inputs to vestibulospinal neurons had remarkably stereotyped amplitudes^20^. Further, Liu et. al. performed genetic loss-of-function experiments that suggest a dominant role for utricular inputs in driving vestibulospinal responses to translation^20^. Follow-up experiments complement these loss-of-function findings with hemibrain EM datasets that establish synaptic connectivity between ipsilateral otolithic afferents and vestibulospinal neurons^21,22^. To date, the electrophysiological and loss-of-function findings have neither been replicated nor extended.

Vertebrates use vestibular sensory organs in each of two ears to detect imbalance. Vestibulospinal neurons could therefore receive unilateral and/or bilateral input, and this input could be excitatory or inhibitory. Comparing information across ears is key to proper vestibular behavior^26–28^, as revealed following unilateral loss of VIII^th^ nerve input^29,30^. In particular, contralateral inhibition of broad origin^31^, or restricted to the utricle^32,33^, has been proposed as a way to increase sensitivity of central vestibular neurons. Intriguingly, there is an existing anatomical divide between lower (e.g. *Hyperoartia)* and higher (e.g. *Mammalia*) vertebrates regarding the lateralization of excitatory and inhibitory input^2^. To date, only ipsilateral excitatory input has been characterized in larval zebrafish vestibulospinal neurons^20^ leaving open questions of vestibulospinal circuit homology and function.

In this paper, we investigated the nature and origin of synaptic input onto vestibulospinal neurons in larval zebrafish. We began by validating and extending three key findings. First, we observed that vestibulospinal neurons, while silent at rest, can fire sustained trains of action potentials. Next, we used both acute and chronic loss-of-function approaches to establish that phasic responses to translation in the dark originate with the utricle. We then used voltage-clamp recordings to characterize the spontaneous excitatory (as done before) and inhibitory (novel) synaptic inputs. While excitatory synaptic inputs on vestibulospinal neurons were separable into discrete event amplitudes, in most cases these failed a refractory period test. We then used unilateral lesions to map the organization of spontaneous synaptic inputs to vestibulospinal neurons. We found that, like mammalian central vestibular circuits, high amplitude excitatory inputs derive from the ipsilateral ear, whereas inhibitory inputs originate from both the ipsilateral and contralateral ear. Taken together, our work builds on and extends previous findings to characterize the synaptic inputs to vestibulospinal neurons in the larval zebrafish. The similarities we observe to mammalian architecture and ability to replicate basic findings across labs solidify the utility of the larval zebrafish to understand the synaptic computations that mediate vestibulospinal reflexes so crucial for vertebrate balance.

## RESULTS

### Vestibulospinal neurons encode body translation using utricular sensory inputs

We first characterized the basic electrophysiolgical properties of larval zebrafish vestibulospinal neurons. We performed *in vivo* whole cell patch clamp recordings in the dark in fish that were 4-12 days post fertilization (dpf, n=21 cells, Figure 1A). Dye in the recording solution allowed *post-hoc* confirmation of vestibulospinal identity by visualization of descending axons. Neurons had high input resistance (236±130 MΩ) and resting membrane potential of −67±5 mV (Table 1). At rest, approximately half (12/21) of vestibulospinal neurons showed no spontaneous action potentials (Figure 1B, top); the remaining 9 had an median firing rate at rest of 0.9 Hz (range: 0.02-16.3 Hz). Silent vestibulospinal neurons could however sustain a high tonic firing rate (90.3±68.5 Hz) following current injection (largest depolarizing step per cell: 84-322 pA) (Figure 1B, bottom). The high rheobase of 111.8±78.9 pA (Figure 1C) in cells that were silent at rest suggests the lack of spontaneous firing activity reflects a high spiking threshold in these neurons. All action potentials had a mature waveform with a median spike amplitude of 50.9 mV (Figures 1D and 1E). We conclude that, similar to *Xenopus* vestibular neurons^34,35^, larval zebrafish vestibulospinal neurons are largely silent at rest but capable of firing sustained trains of action potentials.

**Figure 1:**
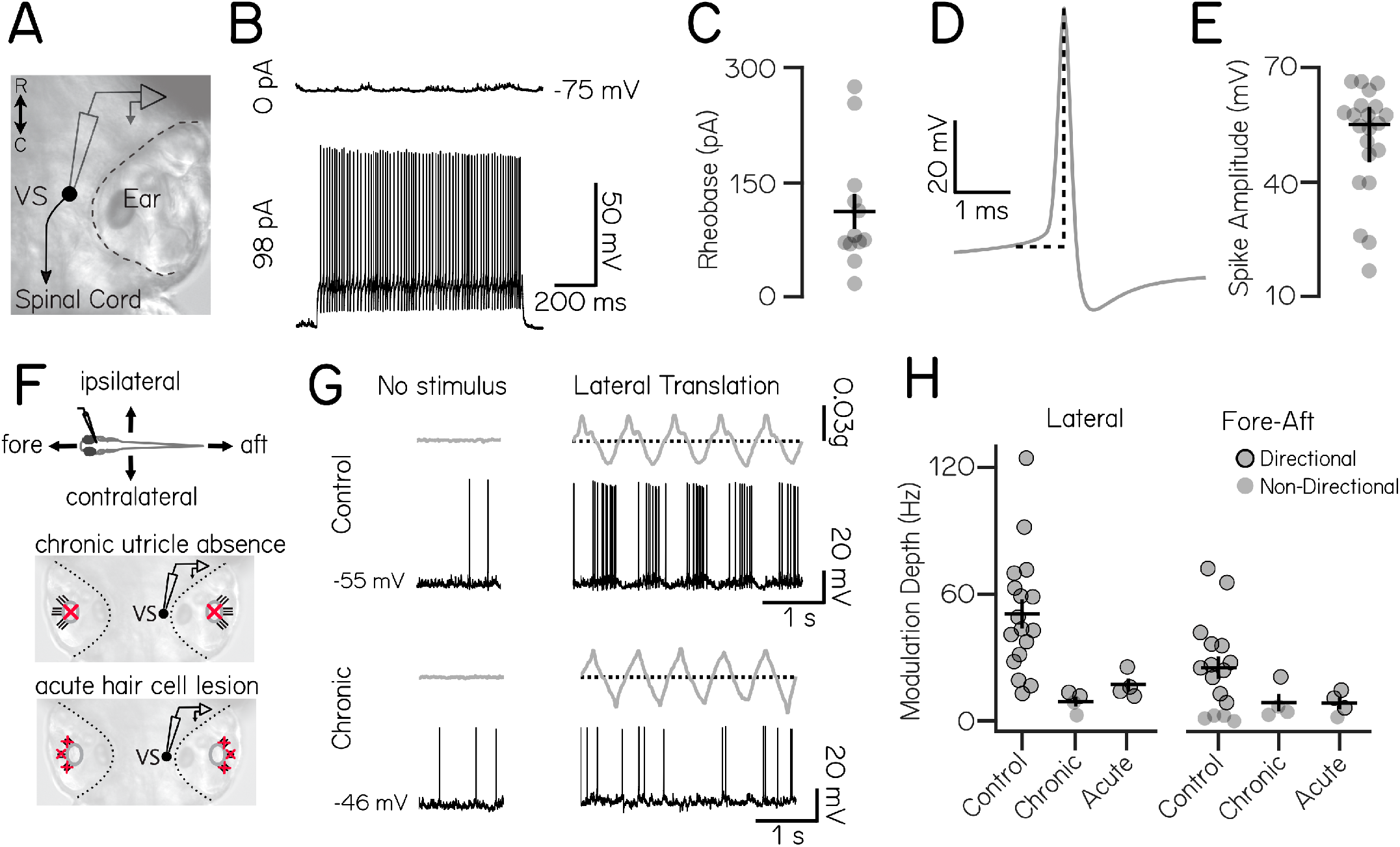
Vestibulospinal neurons encode utricle-derived body translation. (A) Schematic of spinal-projecting vestibulospinal (VS) neuron targeted for electrophysiology (B) Vestibulospinal membrane potential at rest (top, 0 pA) and in response to current injection (bottom, 98 pA) (C) Rheobase (mean±SEM) across 12 non-spontaneously active cells (D) Example action potential waveform with amplitude (dotted line) (E) Action potential average amplitude (median±IQR) across 21 cells (F) Immobilized fish were manually translated in the fore-aft or lateral axes (top). Vestibulospinal neurons were recorded in control and after two manipulations: first, in *otogelin* mutants (middle) that do not develop utricles (red “x”) and second, after chemically-induced hair cell (red “x”) death (bottom) (G) Accelerometer (gray) and voltage trace (black) from a neuron in a control fish (top) showing action potentials in phase with translation. In contrast, activity from a *otogelin* mutant is unaligned with translation (H) Modulation depth of spiking response (mean±SEM) is disrupted in both the lateral (left) and fore-aft (right) direction after both chronic and acute disruption of the utricle. Gray circles are neurons, black outlined circles denotes statistically significant directional responses.

**Table 1:** Spontaneous properties across conditions.

|  | Unit | Control | Cs Control | Ipsi Lesion | Contra Lesion | Bilateral Lesion | *rock solo* |
| --- | --- | --- | --- | --- | --- | --- | --- |
| Resting Membrane Potential | mV | -67.3 | -60.8 | -63.7 | -63.8 | -68.9 | -66 |
| Series Resistance | MΩ | 34.7 | 25.8 | 43.6 | 45.7 | 38.4 | 31.2 |
| Input Resistance | MΩ | 236 | 202 | 264 | 178 | 310 | 228 |
| Resting Firing Frequency | Hz | 1.4 | - | - | - | 0 | 0 |
| Rheobase | pA | 111.8 | - | - | - | 38.9 | 58.8 |
| Resting EPSC Frequency | Hz | 101.3 | 65.4 | 30.2 | 59.2 | 46.4 | 119.2 |

We next assayed the responses of vestibulospinal neurons to sensory stimulation. We provided an oscillatory translation – in the dark – either along the fore/aft or lateral axis of the fish during whole-cell recordings (Figure 1F, top). To ensure that we did not miss evoked responses due to insufficient sensory stimulation, when recording from neurons that were completely silent at rest (n=9/17 cells) we injected a small bias current (91.9 ± 79.6 pA), following previously published methods^20^. The presence or absence of this depolarizing current did not affect further conclusions and so all cells were combined for analysis.

We observed that vestibulospinal neurons fired phasically during translation (Figure 1G, top, response to example lateral translation). In both lateral and fore-aft directions, the majority of cells had a single peak in firing rate during an oscillation cycle and a single anti-phase firing pause (“simple” tuning; 15/17 lateral; 10/16 fore-aft), but a small percent of cells had multiple peak/pauses in firing (“complex” tuning; 2/17 lateral; 2/16 fore-aft) as had previously been observed in the fore-aft axis^20^. We quantified a cell’s directional tuning in each translational axis (lateral or fore-aft) by calculating the difference between the peak firing rate and the minimum firing rate during an oscillation cycle (referred to as the “modulation depth”) (Methods)^36^. This method performed best qualitatively for analyzing both simple and complex tuning cells, and was strongly correlated with comparable tuning metrics such as taking the difference between peak phasic response and the 180deg anti-phase response (Pearson’s correlation, ρ=0.99). We then tested for statistically significant directional tuning by comparing a cell’s modulation depth for that stimulus relative to that derived from randomly shuffled data. We subsequently categorized each cell as “directional” or “non-directional” for each axis. All recorded neurons (17/17) responded directionally to lateral stimuli (modulation depth 50.6±28.6 Hz), and most (12/16) were directionally responsive to fore/aft stimuli (modulation depth 25.3±21.7 Hz) (Figure 1H, directional neurons circled). Most neurons were directionally tuned to the peak acceleration of the stimulus towards the contralateral side (6/10) and rostral direction (7/10 neurons). We conclude that activity of vestibulospinal neurons can encode translation.

Previous loss-of-function studies established that the utricle is the dominant source of sensory information about body tilts in larval zebrafish^6,14,16^. We next asked if the evoked responses we observed reflected activity originating in the utricular macula. We adopted a loss-of-function approach, recording from vestibulospinal neurons in *otogelin* mutants that fail to develop utricles (Figure 1F, chronic utricle absence)^37^. Neurons in mutant fish could still fire action potentials, but failed to respond phasically (Figure 1G, bottom). Modulation depth was decreased in mutant fish compared to controls in both the lateral (9.2±4.8 Hz) and fore-aft axis (8.7±8.3 Hz). Among recordings from mutants (n=4) two neurons met our criteria as “directionally responsive” for lateral translation and one neuron was directionally responsive for fore/aft translation, but modulation depth was low in both directions (Figure 1H, black circles). This data suggest that the bulk of directionally-sensitive inputs to vestibulospinal neurons originates from the utricle.

To control for possible compensatory mechanisms in *otogelin* mutants, we also measured vestibulospinal neuron responses to translation after acute chemo-ablation of inner-ear hair cells (Figure 1F, acute hair cell lesion). Similar to the *otogelin* mutants, after acute chemo-ablation, modulation depth was reduced dramatically in both the lateral (16.8±6.1 Hz) and fore-aft axes (8.4±5.6 Hz) (Figure 1H). Across both acute and chronic *(otogelin)* utricle manipulations, modulation depth was strongly affected by lesion condition, but not stimulus direction (Two-Way ANOVA, main effect of lesion condition F_2,43_=8.5, p=0.0008; main effect of stimulus direction F_1,43_=2.1, p=0.16; interaction effect of lesion condition and stimulus direction F_2,43_ = 1.6, p=0.22) with lower modulation depth in chronic (Tukey’s posthoc test, p=0.004) and acute conditions (Tukey’s posthoc test, p=0.014) compared to controls. Acute lesions did not decrease the fraction of neurons directionally responsive to lateral and fore-aft stimuli (100% lateral directional, 75%fore-aft directional, n=4), but the strength of tuning among responsive cells was low (Figure 1H, directional neurons circled). Collectively, our loss-of-function experiments support the conclusion that utricular input is required for normal phasic responses to translation in vestibulospinal neurons.

Taken together, our data supports earlier findings20 that larval zebrafish vestibulospinal neuron activity reflects sensed destabilization originating with the utricle.

### Larval zebrafish vestibulospinal neurons receive dense spontaneous excitatory and inhibitory synaptic input

We next characterized the complement of excitatory synaptic inputs to vestibulospinal neurons at rest. Neurons received dense synaptic excitatory post-synaptic currents EPSCs (Figure 2A) with a median frequency of 89.4±41.7 Hz (n=35 neurons). EPSCs showed a wide range of amplitudes (median range 136.0 pA). Amplitude distributions were multimodal in all cells, with distinct peaks visible in a probability distribution (Figure 2B). To characterize these peaks, we assigned EPSCs to amplitude ranges that encompassed each peak in the distribution (Figure 2B, line colors). These bins remained stable over time (Figure 2C). Across our data, neurons had a mean of 3.3±1.0 distinct bins (range of 2-5 bins) (115 bins from 35 cells), with a median event amplitude per bin of 39.5±27.0 pA and a median event frequency per bin of 19.2±20.3 Hz. Bin amplitude and frequency were inversely related (Figure 2D).

**Figure 2:**
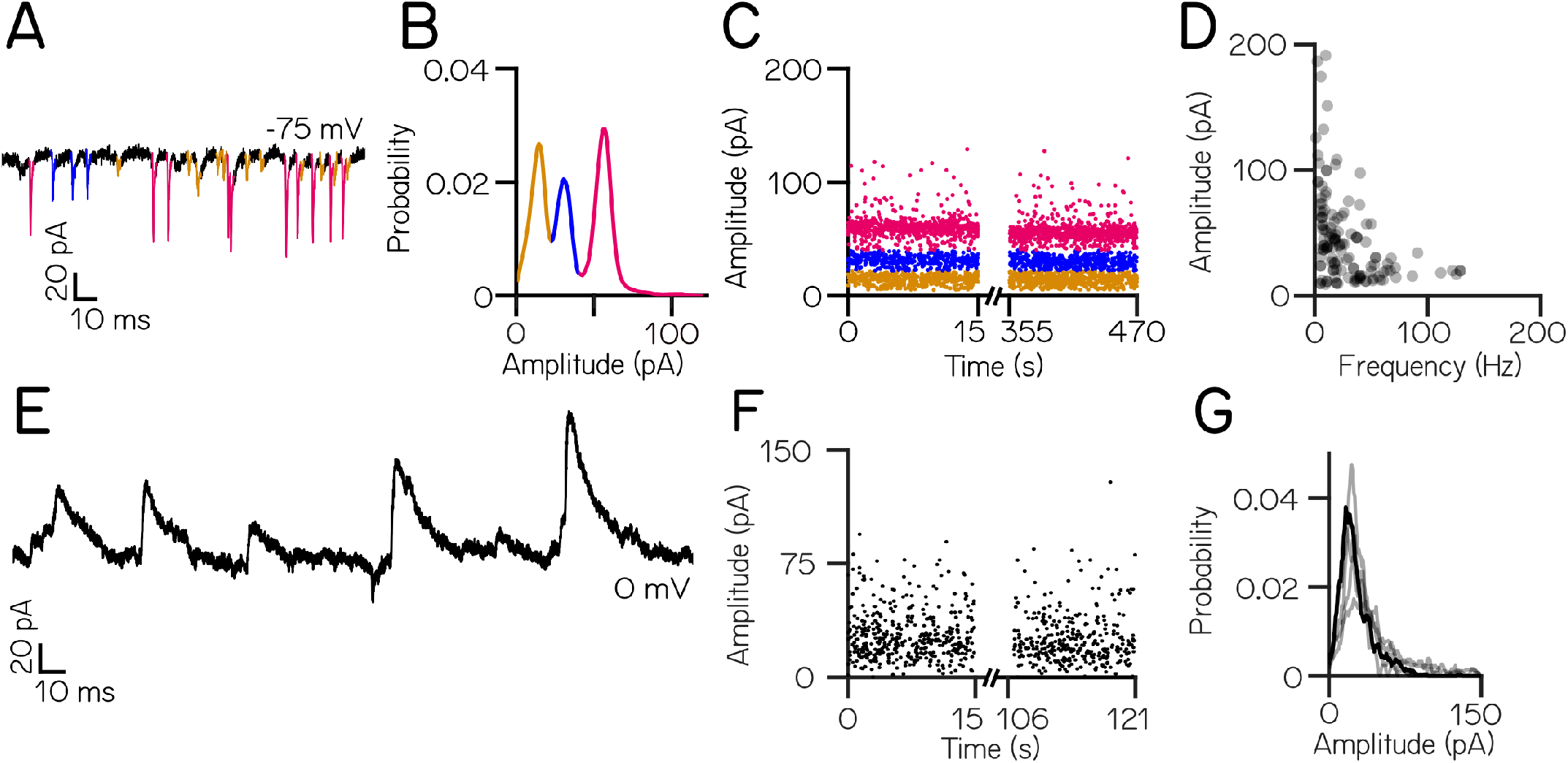
Larval zebrafish vestibulospinal neurons receive dense spontaneous synaptic input. (A) Distinct amplitudes (color) in excitatory post-synaptic spontaneous currents (EPSCs) from a neuron held at −75mV (B) EPSC amplitudes from a single vestibulospinal neuron show 3 distinct probability peaks, or “bins” (color) (C) EPSC bins are stationary in time (D) EPSC amplitude as a function of frequency for all bins in all vestibulospinal neurons (115 bins, from 35 cells) (E) Representative current trace from a vestibulospinal neuron held at 0 mV (F) Inhibitory post-synaptic current (IPSC) amplitudes over time (G) IPSC amplitudes for the example neuron in E/F(black line) and other neurons (gray lines) do not show multiple peaks (n=5)

Next, we performed a separate set of voltage-clamp experiments to isolate inhibitory post-synaptic currents (IPSCs). Neurons received spontaneous IPSCs (Figure 2E) with a mean frequency of 22.4±7.4 Hz (range 13.8-32.5 Hz) and a mean amplitude of 29.1±7.3 pA(range 22.5-39.2 pA) (n=5 neurons). Unlike excitatory inputs, spontaneous inhibitory currents did not exhibit distinct event amplitude peaks (Figures 2F and 2G). We conclude that vestibulospinal neurons receive dense excitatory and inhibitory input at rest.

### EPSC events within the same amplitude bin reflect multiple neuronal inputs

Distinct EPSC bins might reflect input from single VIII^th^ nerve afferents with different stable resting amplitudes^24,25^. A previous report reached this conclusion based on comparable recordings done in the light^20^. To test if EPSCs within a distinct amplitude bin in our recordings (Figure 3A) derived from a single afferent neuron (a “unitary” origin), we applied a refractory period criteria to identify EPSC events that occurred within 1 ms of each other (Figure 3B). We reasoned that if EPSC amplitude bins reflect single afferent inputs, there ought be no such examples of refractory period violations from within-bin EPSCs (Figure 3B, left).

**Figure 3:**
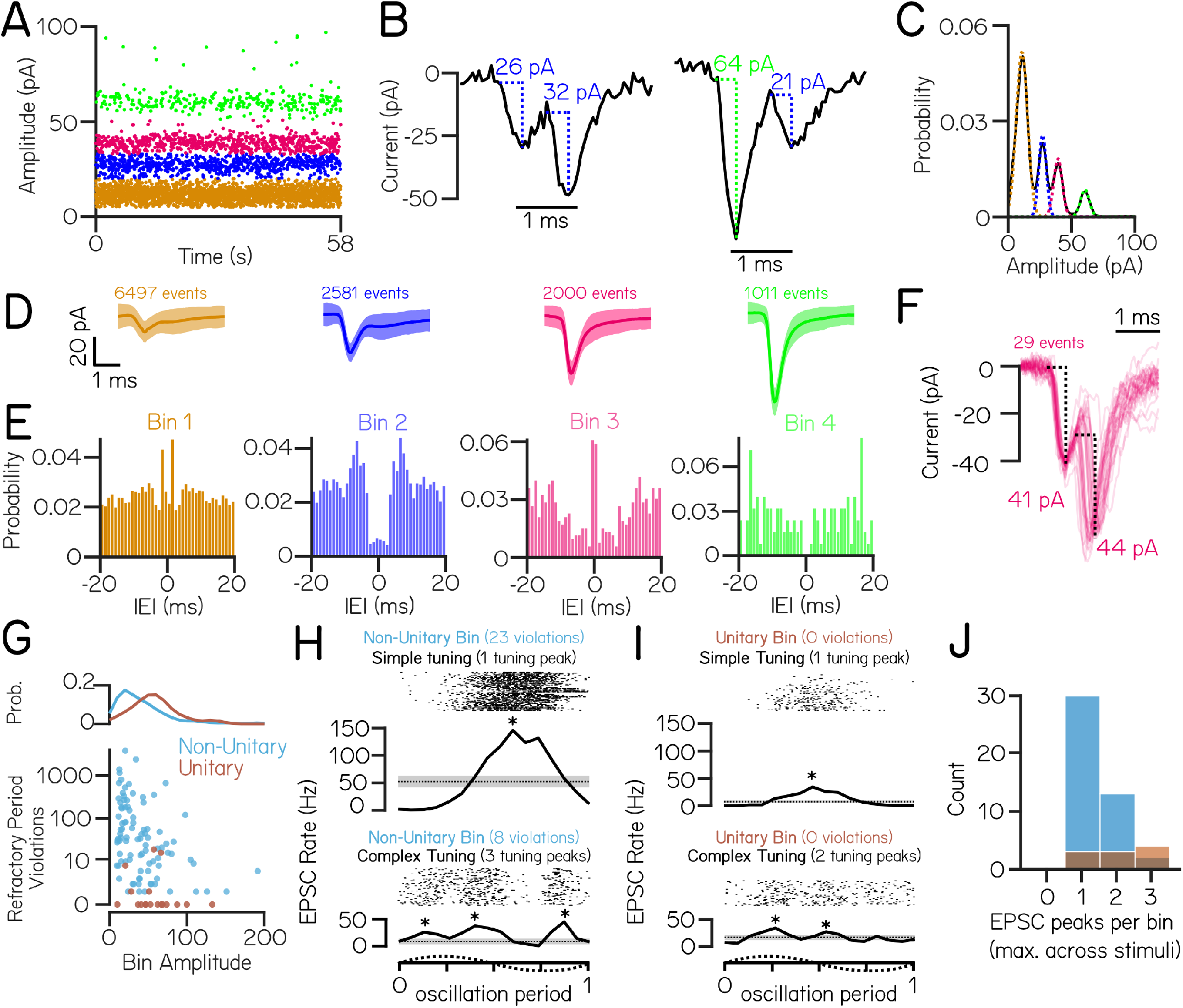
EPSC events within the same amplitude bin predominantly reflect multiple neuronal inputs. (A) An example cell with four stable and discrete amplitude bins (colored by bin) (B) Example EPSC traces demonstrate events that co-occur within 1 ms. Event pairs are either within-bin (left) or across-bin (right) (C) To estimate an upper limit on the expected refractory period violations due to bin overlap, EPSC amplitude distributions were modeled as a sum of individual Gaussians (dashed colored lines) (D) Average waveforms from each EPSC bin (± S.D.) (n=6497, 2581, 2000, & 1011 events per bin) (E) Auto-correlograms show structure of inter-event intervals within an EPSC bin; note peaks near zero in bins 1&3, a non-zero valley for bin 2, and a true valley for the high-amplitude bin 4 (F) Waveforms from EPSC pairs within bin 3 with latencies <1 ms (n=29 event pairs). The large jitter between peaks are inconsistent with the expected profile of an electrochemical synapse. (G) Observed within-bin refractory period violations as a function of bin amplitude for bins assigned as having non-unitary (blue) or unitary (brown) origin. Probability distribution of bin amplitudes shown above. (H) EPSC event timing from two example non-unitary or (I) unitary bins aligned to one oscillation of lateral translation. EPSCs within a single bin can exhibit simple tuning with a single peak in EPSC rate (top), or can have complex tuning with multiple peaks in EPSC rate during oscillation (bottom). Asterisks indicate EPSC tuning peaks. (J) Histogram of the number of EPSC rate peaks per amplitude bin during translation (maximum per bin across lateral and fore-aft stimuli) for non-unitary and unitary EPSC bins.

To account for sources of error (e.g. noisy bin assignations) that might lead us to over-estimate within-bin refractory period violations in our data, we compared empirical within-bin refractory period violations to a model estimate of expected violations. Briefly, each EPSC amplitude bin was modeled as a Gaussian distribution centered around the amplitude of each bin, and an upper limit on expected within-bin refractory period violations was set by the overlap in these modeled Gaussians and the frequency of event in each bin (Methods, Figure 3C). We accounted for the possibility that some trial recordings without refractory period violations were too brief to detect violations in low-frequency EPSC bins, and excluded any such bins from further analysis (4/115 bins).

We first examined EPSC refractory period violations in an example cell with 4 amplitude bins (Figures 3A and 3C). Refractory period violations were a minority of events within each bin, evident in the stereotyped shape of the average waveform of EPSCs (Figure 3D). Only one amplitude bin (Bin 4, green) had no refractory period violations, as expected if bins had a unitary origin. Auto-correlograms of EPSC inter-event intervals within an amplitude bin showed that this potentially unitary bin had a distinct valley near 0 ms (Figure 3E), consistent with previous work^20^. The three remaining bins had more within-bin refractory period violations than estimated by our model of bin overlap, consistent with a non-unitary origin.

We noticed that two bins (bins 1 & 3) exhibited significantly more within-bin refractory period violations than expected based on event frequency alone, reflected by a peak near 0 ms in their auto-correlograms (Figure 3E); these high-violation bins were not uncommon across bins from all cells (24/111 bins). These bins might reflect the presence of compound events (i.e. mixed electrical and chemical synapses) deriving from the same afferent input. Evidence for mixed synapses between the larval zebrafish VIII^th^ nerve comes from electrophysiology/pharmacology and electron microscopy20 in larval zebrafish, from electrophysiology and electron microscopy in other teleosts^38^, and from immunofluorescence and electron microscopy in the rat^39^. However, waveforms of high-probability compound events did not have the expected shap eof a classical electrochemical synapse with an electrical event followed by a stereotyped low-jitter chemical event (Figure 3F). Additionally electrochemical synapses typically consist of a high amplitude electrical event followed by a smaller amplitude chemical event^20,40^, not two events of comparable size as observed here. Finally, these event pairs – while more common than expected by chance – composed a very small percent of the total events within an amplitude bin (29/2000 events in bin 3). As we did not directly test whether events originated from electrical/chemical origins, our data do not speak to whether vestibulospinal neurons generally receive input from mixed synapses. Nevertheless, we conclude it is unlikely that the presence of mixed synapses caused us to dramatically underestimate the presence of unitary EPSC bins.

We then turned to see if our findings generalized across amplitude bins over all cells. As in our example cell, the majority (93/111) of amplitude bins had more violations than expected if they originated from a single afferent unit (“non-unitary”, Figure 3G). Compared to non-unitary bins, the few bins that passed our refractory test consisted of higher amplitude (median 54.9 pA unitary vs. 31.1 pA non-unitary, Figure 3G) and lower frequency EPSCs (median 9.7 Hz unitary vs. 21.6 Hz non-unitary). Vestibular afferents are commonly classified with respect to the stereotypy of their inter-spike intervals, falling into one of two classes: regular or irregular^41^. The inter-event intervals of the putative unitary bins were consistent with irregular afferent input (median coefficient of variation = 0.93±0.08).

To further test whether EPSC amplitude bins were consistent with a unitary origin, we investigated the sensory tuning of EPSCs within an amplitude bin using oscillatory translations of the fish during whole-cell recordings. EPSCs within an amplitude bin could either be tuned similarly, with a single peak in EPSC rate (“simple” tuning) or tuned disparately, with multiple peaks in EPSC rates during an oscillation (“complex” tuning). As vestibular afferents only respond in a single phase direction^42–44^, we reasoned that EPSC bins that exhibit multiple peaks in EPSC rate during an oscillation must originate from multiple afferents with disparate directional tuning. Conversely, EPSC bins that exhibit simple tuning to the translation stimulus could either be derived from a single afferent or from multiple converging afferents with the same preferred stimulus direction.

EPSCs from amplitude bins that were determined to be non-unitary by refractory period violations had examples of both simple (30/45 bins) and complex (15/45 bins) tuning to translation (Figures 3H and 3J), which is consistent with the hypothesis that these EPSCs derive from multiple afferent inputs. Surprisingly, among the EPSC bins determined as putatively unitary by refractory period violations, we still identified bins that had simple (3/10) and complex (7/10) EPSC tuning (Figures 3I and 3J). This result strongly suggests that the majority of EPSC amplitude bins originate from multiple afferent sources with only 5% of bins being consistent with a single afferent source (no refractory period violations and simple EPSC tuning). We conclude that in our recordings nearly all bins are comprised of multiple inputs, but a handful of high-amplitude, low-frequency event bins may be consistent with input from single irregularly-firing VIII^th^ nerve afferents.

### High-amplitude excitatory synaptic inputs originate from ipsilateral ear

Our loss-of-function experiments suggest that sensory-driven input to vestibulospinal neurons is predominantly utricular. As each ear contains a utricle, inputs to a given neuron could originate from ipsilateral or contralateral utricular afferents. To differentiate ipsilateral and contralateral contributions we performed voltage-clamp recordings of spontaneous EPSC activity in vestibulospinal neurons after removing the utricle either ipsilateral or contralateral to the recorded neuron (Figures 4A and 4B). We found that the number of EPSC amplitude bins per cell differed across lesion conditions (Kruskal-Wallis H(2)=10.2, p=0.006) (Figure 4C). After ipsilateral lesion, neurons had fewer EPSC bins (median 1 vs 3; n=9 lesion, n=5 control; Dunn-Sidak post-hoc test, p=0.006). In contrast there was no change after contralateral lesion (median 2.5 bins; n=6; Dunn-Sidak post-hoc test, p=0.54). Further, the amplitude of EPSC bins also differed across conditions (Kruskal-Wallis H(2)=11.7, p=0.003) (Figure 4D). EPSC bins after ipsilateral lesion were lower amplitude than controls (median 11.4 pA vs. 43.2 pA; Dunn-Sidak post-hoc test, p=0.002), but contralateral lesions did not affect EPSC bin amplitudes (median 25.4 pA; Dunn-Sidak post-hoc test, p=0.53). EPSC bin frequency was not changed across lesion conditions (Kruskal-Wallis H(2)=0.06, p=0.97) (Figure 4E). We conclude that high-amplitude, low-frequency EPSCs derive from ipsilateral inputs (Figure 4F). In contrast, lower-amplitude EPSCs persist after both ipsilateral and contralateral lesions, which might reflect either an extra-vestibular origin or an incomplete lesion.

**Figure 4:**
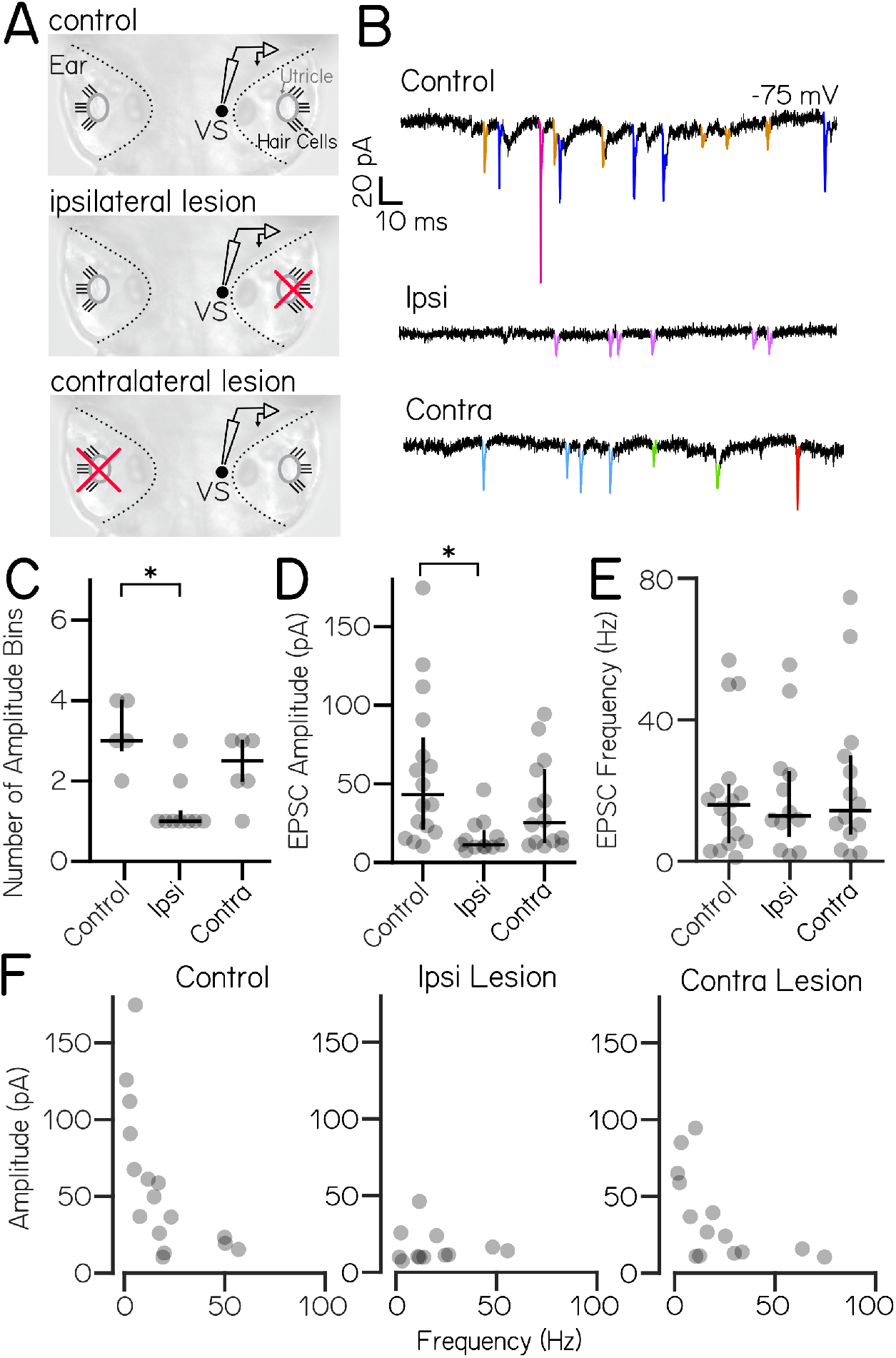
High amplitude spontaneous excitatory inputs originate in the ipsilateral ear. (A) Lesion schematic: The utricle (gray circle) was physically removed (red “x”) either ipsilateral or contralateral to the recorded vestibulospinal neuron (black circle, “VS”) (B) Example current traces from neurons held at −75 mV from control (top), ipsilateral (middle), and contralateral (bottom) experiments; EPSCs in color (C) Number of EPSC amplitude bins per cell (median ± IQR in black) is decreased after ipsilateral, but not contralateral lesion (D) EPSC bin amplitudes (gray circles, median ± IQR in black) are decreased after ipsilateral, but not contralateral, lesion (E) Frequency of events in EPSC bins (gray circles, median ± IQR in black) is unchanged after ipsilateral or contralateral lesion compared to control cells (F) EPSC amplitude vs. frequency for each bin (gray circles) in control and after ipsilateral/contralateral lesions. High amplitude bins are lost after ipsilateral lesion

### Inhibitory inputs originate with both ipsilateral and contralateral ears

We then asked whether inhibitory synaptic input onto vestibulospinal neurons originated from the ipsilateral or contralateral ear. We quantified spontaneous IPSCs after ipsilateral or contralateral utricular lesions. In control cells without peripheral lesions, spontaneous IPSCs onto vestibulospinal neurons occurred at a frequency ranging from 13.7-32.5 Hz (n=5 cells). After ipsilateral lesion, we found that the range of IPSC frequencies increased (1.1-28.5 Hz), where half of the recorded cells had IPSC frequency that dropped markedly compared to controls (n=4/8 cells falling below 10 Hz). In contrast, IPSC frequency was comparable to control cells in the other half of ipsilateral lesion cells (Figures 5A and 5B). Interestingly, we found that contralateral utricular lesions had a similar effect on the frequency of IPSC input to vestibulospinal neurons. After contralateral utricular lesions, IPSC frequency range increased compared to control cells (0.8-34 Hz). A fraction of cells experienced a drastic reduction in IPSC frequency compared to controls (n=2/6 cells falling below 10 Hz), while the remaining cells had comparable IPSC frequency to control cells (Figures 5A and 5B). Ipsilateral and contralateral lesions did not affect the amplitude of remaining IPSCs in any of the cells (One-Way ANOVA F(2,16)=0.15, p=0.86) (Figure 5C). Our data suggest that spontaneous IPSCs onto vestibulospinal neurons can reflect vestibular input of utricular origin. Furthermore, our data are consistent with a model where an individual vestibulospinal neuron receives the majority of its inhibition from either the ipsilateral or contralateral ear, rather than a convergence from both ears.

**Figure 5:**
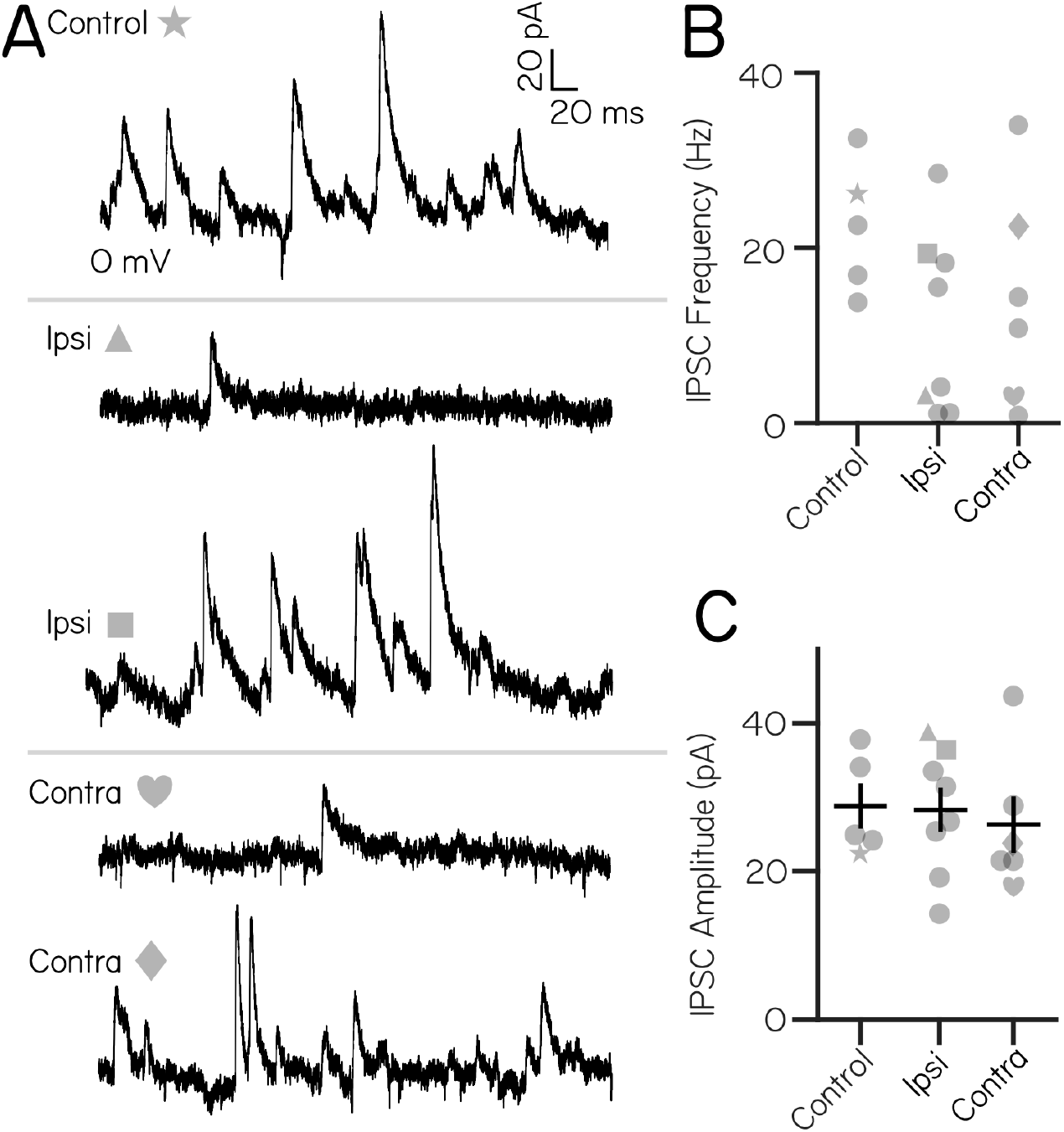
Inhibitory current inputs have ipsilateral and contralateral vestibular sensory origins. (A) Control trace from a neuron held at 0mV shows inhibitory input at rest (top). After ipsilateral (middle) or contralateral (bottom) utricular lesion, some cells experience strong loss of inhibitory currents, while others appear unaffected. (B) Distribution of IPSC frequency after ipsilateral or contralateral utricular lesions. Symbols correspond with example neurons in panel (A). No control neuron (n=5) experienced IPSC frequency less than 10 Hz. After ipsilateral lesions, 4/8 neurons experienced IPSC frequencies less than 10 Hz. After contralateral lesion, 2/6 neurons received IPSCs below 10 Hz. (C) IPSC amplitude is unchanged across control and lesion conditions (mean±SEM)

## DISCUSSION

Vestibulospinal neurons are part of a critical and evolutionarily-ancient circuit that transforms peripheral sensations of imbalance into postural motor reflexes. Here, we used the larval zebrafish to investigate the source and structure of excitatory and inhibitory synaptic inputs onto zebrafish vestibulospinal neurons. We began by confirming and extending findings from a previous report^20^. We confirmed that neurons were silent at rest yet capable of finding sustained trains of action potentials upon depolarization, and that neurons responded systematically to oscillatory translation in the dark. We replicated the observation that genetic loss of utricular function disrupted these vestibular responses, and extended this finding to acute bilateral lesions of the ear. We confirmed that vestibulospinal neurons received excitatory synaptic inputs of characteristic amplitude. However, in our recordings, the bulk of these characteristic amplitude bins failed a refractory period test suggesting a non-unitary origin. We discovered that vestibulospinal neurons also receive strong inhibitory inputs. Finally, we used acute unilateral lesions to show that loss of ipsilateral input disrupted the highest amplitude excitatory inputs, and that both ipsilateral and contralateral lesions could disrupt IPSCs. Together, our work both validates a recent characterization of vestibulospinal neurons, and extends that work to map circuit-level inputs to larval zebrafish vestibulospinal neurons.

Linear encoding at central vestibular synapses is thought to be important for encoding of head/body position. One way to achieve linear encoding by the maintenance of stable, frequency-invariant EPSC amplitudes^24^. Stable EPSC amplitudes can be instantiated by a number of pre- and post-synaptic molecular mechanisms that keep overall charge transfer across the synapse the same over time^25^. A single afferent should therefore have stable excitatory drive over time. If the stable amplitudes of each afferent are different from each other, then inputs onto a postsynaptic neuron should be separable by EPSC amplitude, as was previously reported^20^. We observe that EPSCs onto vestibulospinal neurons fall into discrete amplitude bins that are stable across time/trials.

However, our data is largely inconsistent with the model that EPSCs within a bin reflect a singular afferent input. Instead, we suggest that a bin consists of input from several afferents with roughly the same stable EPSC amplitude. Each individual afferent maintains stable charge transfer over time, as proposed^24,25^. In this model, the ability to differentiate single afferent inputs while recording post-synaptically is limited by (1) the intrinsic noise of our recordings and (2) the number of inputs converging onto the post-synaptic cell. As our intrinsic noise was low, our preparation likely resulted in more spontaneous and sensory-evoked inputs compared to previous preparations. Importantly, our data nevertheless support a model where afferent synapses onto vestibulospinal neurons achieve linear encoding of head and body movement through stable excitatory drive.

The only major difference between the experimental preparations here and in20 was that our recordings were performed exclusively in the dark while theirs were in ambient light. Notably, the previous report focused on analysis of a subset ( 50%) of recorded vestibulospinal neurons that had one or more amplitude bins whose activity was consistent with a singular origin (M. Bagnall, personal communication). We hypothesize that differences might reflect visual or state-dependent modulation of presynaptic inputs to vestibulospinal neurons. Both visual input1 and behavioral state45 can profoundly impact vestibular neuron activity, and presynaptic inputs to vestibular neurons are thought to be the site for visually-guided motor learning^46^. In larval zebrafish, such modulation could originate from visually-responsive47 dopaminergic neurons that when activated drive vestibulospinal neuron activity^48^. We therefore build on and expand models of vestibulospinal circuit organization to offer a tractable way to understand – at a synaptic level – how extra-vestibular information influences sensed imbalance.

Electrophysiological studies have established a basic map of excitatory and inhibitory vestibular synaptic inputs in a number of vertebrate species (Figure 6). Vestibulospinal neurons in all species are defined, in part, by receiving excitatory input from the ipsilateral vestibular nerve. There is an existing divide, however, between “lower” and “higher” vertebrates regarding the role of contralateral vestibular input. Studies in mammals and frogs have shown evidence of contralateral inhibition^27,32,33,49–51^, which has not been reported in studies of other teleost fish or non-jawed vertebrates^38,52^; conversely, vestibulospinal neurons in many non-mammalian vertebrates receive contralateral excitation^38,51,53^, which is not as commonly seen in mammalian cells^33,49,50,54^ (Figure 6A). In the cat, where the circuit has been the most carefully mapped, vestibulospinal neurons receive excitatory utricular inputs predominantly from the ipsilateral ear, cross-striolar inhibition from the ipsilateral ear, and commissural inhibition from the contralateral ear^33,54–57^. The source of inhibitory inputs has been of particular interest when discussing circuit function in mammals, as commissural and cross-striolar inhibition are thought to increase the sensitivity of central vestibular neurons to sensory stimuli^31,58^ and to play a role in vestibular compensation^59,60^.

**Figure 6:**
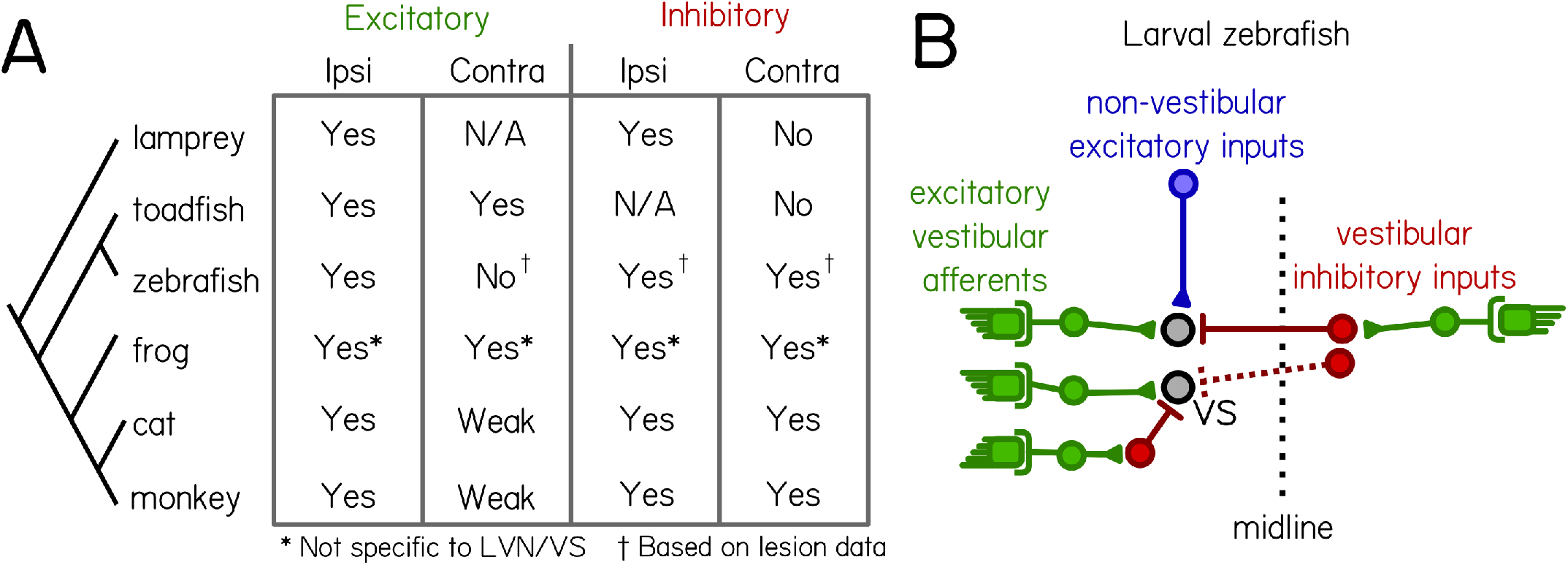
Comparative synaptic architecture of zebrafish vestibulospinal neurons. (A) Summary of previous circuit mapping of functional synaptic connections between vestibular afferents and secondary vestibular neurons across species (Lamprey^52^. Toadfish^38^. Zebrafish^20^ and current study. Frog^27,51,53,65^. Cat^33,54–57^. Monkey^49,50^). All characterizations were from vestibulospinal neuron homologues, except for the frog (asterisk) where data was not specific to vestibulospinal neurons in the lateral vestibular nucleus. Connections were determined by afferent activation, except where only afferent lesion data from the current study was available (dagger). (B) vestibulospinal neurons receive convergent high amplitude excitatory inputs (green) from irregular afferents originating with the ipsilateral utricle (see also^20^), low-amplitude excitatory inputs (blue) from extra-vestibular sources and inhibitory inputs (red) from either the ipsilateral or contralateral utricle.

Our data suggest that individual zebrafish vestibulospinal neurons receive: 1) high-amplitude excitatory inputs exclusively from the ipsilateral utricle, 2) utricle-independent low-amplitude excitatory inputs, 3) inhibitory inputs primarily from either the ipsilateral or contralateral utricle, but likely not both (Figure 6B, right). We do not see a change to spontaneous excitatory inputs after contralateral lesions. However, this may reflect the limits of our loss-of-function approach; future experiments could use afferent stimulation to definitively address whether excitatory inputs originate from the contralateral VIII^th^ nerve. We therefore conclude that zebrafish larvae are closer in their circuit organization to mammals than to other lower vertebrates, based on the presence of contralateral inhibition and the lack of appreciable contralateral excitation. As the larval zebrafish has grown increasingly popular as a useful model to study vestibular circuit function, circuit mapping as we present here will be necessary not only for understanding the logic of this sensory circuit, but for comparing how findings in the zebrafish extend to other species. As the larval zebrafish has increasingly been used as a useful model for studying vestibular circuit function^12,20,61^, development^8,62^, and behavior, it is necessary that we establish a replicable consensus for the synaptic connections within vestibular circuits in the fish. Here, we validated and extend our understanding of the nature and origin of synaptic inputs onto central vestibulospinal neurons in the larval zebrafish. Our work is therefore a major a step towards understanding how sensed imbalance is transformed by these conserved neurons into commands to stabilize posture.

## ACKNOWLEDGMENTS

Research was supported by the National Institute on Deafness and Communication Disorders of the National Institutes of Health under award numbers R00DC012775, R56DC016316, R01DC017489 to DS, and F31DC019554 to KRH. The authors would like to thank Başak Sevinç for assistance with fish maintenance and care, and Katherine Nagel, Niels Ringstad, Mike Long, Dan Sanes, Dora Angelaki, Martha Bagnall along with the members of the Schoppik and Nagel lab for their valuable feedback and discussions.

## AUTHOR CONTRIBUTIONS

Conceptualization: KRH and DS, Methodology: KRH, KH, and DS, Investigation: KRH and KH, Visualization: KRH Writing: KRH Editing: DS, Funding Acquisition: KRH and DS, Supervision: DS.

## MATERIALS AND METHODS

### Fish Care

All procedures involving zebrafish larvae *(Danio rerio)* were approved by the Institutional Animal Care and Use Committee of New York University. Fertilized eggs were collected and maintained at 28.5°C on a standard 14/10 hour light/dark cycle. Before 5 dpf, larvae were maintained at densities of 20-50 larvae per petri dish of 10 cm diameter, filled with 25-40 mL E3 with 0.5 ppm methylene blue. After 5 dpf, larvae were maintained at densities under 20 larvae per petri dish and were fed cultured rotifers (Reed Mariculture) daily.

### Fish Lines

Experiments were done on the *mitfa-/-* background to remove pigment. For chronic bilateral utricular lesions, fish with homozygous recessive loss-of-function mutation of the inner ear-restricted gene, *otogelin* (otog-/-), previously called *rock solo*^AN66^ 37 were visually identified by a lack of utricular otoliths.

### Electrophysiology

Larval zebrafish between 3-15 dpf were paralyzed with pancuronium bromide (0.6mg/mL) in external solution (in mM: 134 NaCl, 2.9 KCl, 1.2 MgCl_2_, 10 HEPES, 10 glucose, 2.1 CaCl_2_) until movement ceased. Fish were then mounted dorsal-up in 2% low-melting temperature agarose and a small incision was made in the skin above the cerebellum. Pipettes (7-9 MOhm impedance) were lowered to the plane of the Mauthner cell body, illuminated by infrared (900nm) DIC optics. Putative vestibulospinal neurons were targeted for patching using soma size and proximity to the Mauthner lateral dendrite as guides. Initial targeting conditions were determined with reference to cells labelled by spinal backfills. Subsequent recordings were determined to be from vestibulospinal neurons by post-recording analysis of anatomical morphology, confirming a single ipsilateral descending axon using either widefield fluorescence or confocal microscopy. All electrophysiological measurements were made in the dark. Pipettes were filled with dye (Alexa 647 hydrazide, Thermo Fisher A20502) in the internal solution (in mM: 125 K-gluconate, 3 MgCl_2_, 10 HEPES, 1 EGTA, 4 Na_2_-ATP). For recordings with voltage clamp trials at 0 mV holding potential, pipettes were filled with an internal solution to prevent action potentials (in mM: 122 CsMeSO_3_, 5 QX-314 Cl, 1 TEA-Cl, 3 MgCl_2_, 10 HEPES, 1 EGTA, 4 Na_2_-ATP). During trials to determine rheobase and maximum firing frequency, cells were injected with 3 pulses (0.5 sec pulse duration) of decreasingly hyperpolarizing current, and 7 pulses (0.5 sec pulse duration) of increasingly depolarizing current. The magnitude of current steps were scaled for each cell until spikes were seen during the strongest depolarizing current. The average current step size for control vestibulospinal neurons was 21.75 pA (range 12-46 pA) with average peak depolarizing current of 152.25 pA (range 84-322 pA). In current clamp trials during linear acceleration stimuli, cells were often injected with an offset depolarizing current (range 0-292.7 pA, mean 46.8 pA across conditions; 57.6 pA control, 32.8 pA acute bilateral utricle removal, 23.8 pA chronic bilateral utricle removal) until spontaneous action potentials were seen during the baseline period without translation.

### Linear Translation Stimulus

Fish were mounted dorsally on an air table (Technical Manufacturing Corporation) with handles (McMaster Carr 55025A51) mounted underneath for manual translation. The air table was then pushed back and forth in either the lateral or fore/aft direction to produce persistent table oscillations. Acceleration traces of table oscillations were measured using a 3-axis accelerometer (Sparkfun, ADXL335) mounted to the air table. Table oscillations persisted for an average of 11.69±5.05 seconds, with a mean frequency of 1.54±0.17 Hz and peak acceleration of 0.91±0.21 g across all trials (n = 122 trials (73 lateral; 49 fore-aft)). Lateral translation trials were longer in duration than fore-aft trials (13.02 s lateral vs. 9.71s fore-aft; p=2.98*10^-4^ unpaired t-test), and higher in peak acceleration (1.01g lateral vs 0.75g fore-aft; p=3.02*10^-14^), but did not significantly differ in stimulus frequency from fore-aft stimuli (1.52 Hz lateral vs 1.58 Hz fore-aft; p=0.051).

### Electrophysiology Analysis

Data analysis and modeling were performed using Matlab (MathWorks, Natick, MA, USA). For current clamp recordings, action potentials were identified with reference to a user-defined threshold for each trial. Action potential amplitude was calculated as the difference between peak membrane voltage and a baseline voltage (1 ms prior to event peak). Rheobase for vestibulospinal action potential generation was calculated by fitting a line to firing responses as a function of injected current, limited to all current steps above the minimum current injection which elicited spikes. The linear fit was used to solve for the estimated current necessary for each vestibulospinal neuron to fire at 1 Hz.

For voltage clamp recordings, excitatory (EPSC) and inhibitory (IPSC) events were identified through hand-selection of events with a waveform consisting of an initial sharp amplitude rise and exponential-like decay. Exact event times were further identified by local minima or maxima search around hand-selected event times. EPSC amplitudes were determined by subtracting the minima of the event waveform from a pre-event baseline current (0.4 ms before event minima). For EPSCs, amplitude “bins” were assigned manually using the probability distributions of EPSC amplitudes across all trials from the same cell. EPSCs with amplitudes less than 5 pA were excluded from analyses. IPSC amplitudes were determined by taking the difference between the maxima of the event waveform and a pre-event baseline current (2 ms before event maxima).

### Assessment if EPSCs had a unitary origin

To determine if EPSC bins were derived from a single afferent origin, we calculated the number of within-bin EPSCs that occurred within a 1 ms refractory period. We excluded events that occurred within 0.3 ms of each other from this analysis as manual inspection found that these were usually double-selections of the same synaptic event, rather than two separable events. We only rarely observed vestibulospinal EPSC amplitude bins with zero within-bin violations. To minimize Type II errors from overly strict refractory period violation criteria, we modeled an upper limit on the number of within-bin refractory period violations that we would expect to see from the overlap in EPSC amplitude distributions: The probability distribution of EPSC amplitudes (*I*) of each cell was estimated as as a sum of Gaussian distributions, where the number of distributions was set to the number of EPSC amplitude bins in that cell. Each amplitude bin was fit with 3 free parameters for height (*h*), center (*c*) in pA, and standard deviation (σ):

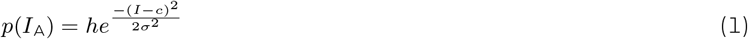

Bin centers (*c*) were constrained to fall within bin amplitude cut-offs. For cells with only one amplitude bin or for the highest amplitude bin, *c* was constrained at the upper bound to be one standard deviation above the peak amplitude probability. Bin heights (*h*) were constrained to be at least half of the maximum value of the empirical probability distribution within the amplitude bin limits. For a given bin of interest (*A*), we used the modeled probability distributions to calculate the number of expected false-positive refractory period violations (an across-bin event pair being falsely counted as a within-bin event pair, *φ_A_*) according to the formula:

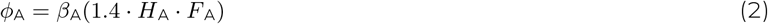

where *β*_A_ is the number of observed EPSC events falling with amplitude bin, *A* (defined by EPSC amplitude thresholds from *a* to *b* pA), *H*_A_ is the hit event rate ((events per millisecond assigned to bin A that truly derived from bin *A*), and *F_A_* is the false-positive event rate for bin A (events per millisecond falsely assigned to bin A when they derived from the overlapping tails of Gaussian distributions from other bins). The joint probability of *H*_A_ and *F*_A_ was multiplied by 1.4 to account for the 0.7 ms window preceding or following any given event to be counted as a refractory period violation (from 0.3 to 1 ms following, or −1 to −0.3 ms preceding). *H*_A_ was calculated using:

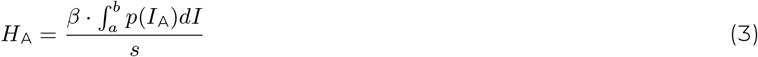

where *β* is the total number of observed EPSCs events across all amplitude bins, and *s* is the trial length in milliseconds. *F*_A_ was calculated using:

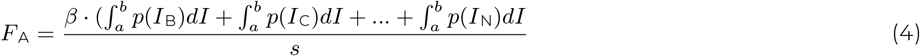

where *p*(*I*_B_) is the estimated probability distribution of the second EPSC amplitude bin, and *p*(*I*_N_) is the n^th^ EPSC amplitude bin.

For each amplitude bin in the cell, φ was calculated, and compared to the number of observed within-bin violations. To determine whether bins with few to no observed refractory period violations occurred solely due to low event frequency, we calculated the number of expected refractory period violations in frequency-matched randomly generated event trains. EPSC bins were classified as unitary afferent bins if the number of empirical within-bin refractory period violations was fewer than the number of expected violations from bin overlap (φ) and if the frequency-matched generated controls had at least 1 observed violation.

### Quantification of sensory responses

Instantaneous spike or EPSC rates were estimated by computing a peri-stimulus time histogram (PSTH) with a time bin width of 1/16 of the oscillation cycle length. A cell’s spiking response (or an amplitude bin’s EPSC response) was defined as the average PSTH across all stimulus cycles in a particular direction (lateral or fore-aft). Modulation depth was determined from this response, defined as the difference between the peak spiking rate and the minimum spiking rate. Cells were considered “directional” if their sensitivity was greater than two standard deviations from the mean modulation depth derived from 100 randomly generated frequency-matched spike trains. The number of firing rate or EPSC rate peaks per oscillation was determined by finding local maxima in the average PSTH. Local maxima were considered a true peak for firing rate if they were greater than two standard deviations from the shuffled mean modulation depth and phase-shifted at least 90° from another true peak. Local maxima were considered a true peak for EPSC rate if they were greater than 1 standard deviation above the mean pre-stimulus baseline EPSC rate for that amplitude bin and phase-shifted at least 90° from another true peak. A cell (for firing rate) or amplitude bin (for EPSC rate) was considered to have “simple” tuning if it had only a single peak in lateral or fore-aft translation, and “complex” if it had more than one peak in a translation direction. For spiking rate, simple/complex tuning was determined separately for lateral and fore-aft directions in cells. For EPSC rate, simple/complex tuning for a single bin was determined by responses to both lateral and fore-aft directions; an amplitude bin that was complexly tuned in either direction was considered complexly tuned overall. EPSC bins that had no significant tuning (0 EPSC peaks) in either direction were excluded from this analysis (n = 1 amplitude bin).

### Utricular Lesions

Chronic bilateral utricular lesions were achieved using mutant larvae *(otogelin)* that do not express *otogelin* and do not develop utricles until 11-12dpf. Acute ipsilateral and contralateral lesions were performed using forceps to rupture the otic capsule and remove the utricular otolith from one ear. This physically removes the sensory organ itself, and likely damages the closely apposed hair cells in the utricular maculae whose spontaneous activity influences vestibular afferents. Acute bilateral lesions were performed through microinjection of 1 mM CuSO_4_ into both otic capsules to kill hair cells^63^, with co-injection of 40 uM FM 1-43 dye (Invitrogen T3163) to label hair cell membranes for visualization. After ipsilateral utricle removal and bilateral copper injection, we saw a marked decrease in spontaneous inputs onto vestibulospinal neurons (data not shown), supporting the hypothesis that these lesions work to impair the firing rate of utricular vestibular afferents. Acute lesions may also impair inner-ear function by diluting the potassium-rich ionic composition of ear endolymph that is critical for hair cell function^64^.

### Statistics

The expected value and variance of data are reported as the mean and the standard deviation or the median and median absolute difference. When data satisfied criteria of normality (Lilliefors test for normality), parametric statistical tests were used, otherwise we used their non-parametric counterparts. Criteria for significance was set at 0.05 and, when applicable, corrected for multiple comparisons.

### Data sharing

All raw data and code for analysis are available at the Open Science Framework

DOI 10.17605/OSF.IO/M8AG9

